# Multiomics study of a heterotardigrade, *Echinisicus testudo*, suggests convergent evolution of anhydrobiosis-related proteins in Tardigrada

**DOI:** 10.1101/2020.10.27.358333

**Authors:** Yumi Murai, Maho Yagi-Utsumi, Masayuki Fujiwara, Masaru Tomita, Koichi Kato, Kazuharu Arakawa

## Abstract

Many limno-terrestrial tardigrades can enter an ametabolic state upon desiccation, in which the animals can withstand extreme environments. To date, studies of the molecular mechanism have predominantly investigated the class Eutardigrada, and that in the Heterotardigrada, remains elusive. To this end, we report a multiomics study of the heterotardigrade *Echiniscus testudo*, which is one of the most desiccation-tolerant species. None of the previously identified tardigrade-specific anhydrobiosis-related genes was conserved, while the loss and expansion of existing pathways were partly shared. Furthermore, we identified two families of novel abundant heat-soluble proteins and the proteins exhibited structural changes from random coil to α-helix as the water content decreased *in vitro*. These characteristics are analogous to those of anhydrobiosis-related protein in eutardigrades, while there is no conservation at the sequence level. Our results suggest that Heterotardigrada have partly shared but distinct anhydrobiosis machinery compared with Eutardigrada, possibly due to convergent evolution within Tardigrada.

## Introduction

Water is an essential universal solvent in living organisms that harbors cellular biochemical reactions. However, many limno-terrestrial tardigrades are able to enter an ametabolic state, known as anhydrobiosis, when they face desiccation, and these organisms can subsequently tolerate almost complete water loss while being able to return to their natural active state when rehydrated ^1-3^. In anhydrobiosis, tardigrades are capable of tolerating many extreme environments, such as high and low temperatures ^4,5^, space vacuum ^6^, high pressure ^7^, and high concentrations of organic solvent ^8^, as well as insults from ultraviolet rays and over 4,000 Gy of gamma radiation ^9-11^.

Early studies on the molecular mechanisms underlying anhydrobiosis initially targeted *Caenorhabditis elegans* ^12^, *Artemia salina* ^13^, and *Polypedilum vanderplanki* ^14^, where the disaccharide trehalose was identified as one of the key compounds, being accumulated at up to 20% of the dry weight during desiccation in *P. vanderplanki*. Trehalose is suggested to protect intracellular components, including membranes and proteins, through water replacement and vitrification ^14,15^. Another of the key factors of the anhydrobiotic machinery is the hydrophilic late embryogenesis abundant (LEA) protein superfamily, which has been identified in both plants and animals ^16^. This widespread protein family exhibits little sequence conservation among groups, but they often show similar structural characteristics, where the protein is mostly natively unfolded in the hydrated state and later folds upon desiccation ^17,18^. Another hallmark is the extreme hydrophilic nature of this protein, which maintains solubility even after ten minutes of heat treatment at 100 °C ^19^. The contribution of LEA proteins in desiccation tolerance is reported to be multifactorial, encompassing protecting membranes, inhibiting the aggregation of proteins, stabilizing cellular components, including vitrified sugar glasses, and serving as a hydration buffer ^20^.

Trehalose and LEA proteins, however, are not the primary machinery of tardigrade anhydrobiosis. Trehalose levels account for only several percent at maximum, and some species even lack measurable amounts of trehalose ^14,21-23^. LEA proteins are observed in the tardigrade transcriptome but are not among the most abundant group of transcripts, suggesting the existence of unique tardigrade components ^24,25^. With the emerging availability of genomic resources ^23,26^, these tardigrade-specific anhydrobiosis-related proteins were first screened using heat solubility assays in the eutardigrade *Ramazzottius varieornatus* ^27,28^. While no sequence similarity to LEA was observed, characteristics of the cytoplasmic abundant heat soluble (CAHS) protein, such as its high abundance, heat solubility, intrinsically unstructured nature, and structural change to alpha helix upon water loss, mirror that of LEA proteins ^27^. These tardigrade-specific anhydrobiosis-related genes initially appeared to be highly conserved among different tardigrades, where the CAHS gene was conserved all across the class eutardigrada in Hypsibiidae, Macrobiotiidae, and Milnesiidae ^23,25,27,29^. Intriguingly, eutardigrades appear to adopt to environments with differing desiccation conditions by adjusting the expression of these genes while maintaining the gene sets. For example, the strong anhydrobiote *R. varieornatus* can immediately enter anhydrobiosis upon desiccation with constitutively high expression of CAHS and other proteins required for anhydrobiosis. On the other hand, semiaquatic *Hypsibius exemplaris* requires 24-48 h of preconditioning before entering anhydrobiosis ^30^ and prepares homologous proteins by de novo expression during this period ^23^, similar to the anhydrobiosis induction method observed in the arthropod *P. vanderplanki* ^31^ and nematode *C. elegans* ^32^. However, conservation in the other class of Heterotardigrada was not observed from transcriptome analysis of a marine species, suggesting the possibility of an alternative set of machineries ^25^.

To date, molecular analysis of Heterotardigrada has been predominantly limited due to the lack of sustainable laboratory rearing systems. Several species of terrestrial heterotardigrades (*i*.*e*., *Echiniscus testudo*) are reported to have comparable or even higher desiccation tolerance than *R. varieornatus* ^33^; therefore, a certain specialized machinery should also exist in these species. Similarly, series of studies have reported the tolerance of heterotardigrades to various extreme environments, including exposure to vacuum ^34^, osmotic pressure ^35^, and high concentrations of environmental toxicant copper ions, which may increase reactive oxygen species ^36^. Evolutionary perspectives on such genetic components would be especially intriguing in Heterotardigrada, since the order Arthrotardigrada predominantly harbors marine species and is considered to be an ancestral group within the phylum Tardigrada ^37^. The existence of LEA proteins in a wide variety of organisms without notable sequence similarity suggests an analogous mechanism in heterotardigrades.

To this end, we conducted a multiomic study of the heterotardigrade *E. testudo* using ultrasensitive methodology ^38-40^ to elucidate the molecular components of anhydrobiosis in this species. This is the first draft genome of a heterotardigrade, in which the tardigrade-unique proteins identified in eutardigrades are not conserved. We identified two families of novel abundant heat-soluble proteins in this genome, which we named *E. testudo* Abundant Heat Soluble (EtAHS), and their structural analogy to the eutardigrade-specific proteins while lacking any sequence similarity suggests convergent evolution of anhydrobiosis machinery in the two classes of tardigrades.

## Results

### Genome of *Echiniscus testudo*

We obtained 33 Gb of DNA sequencing data, corresponding to 85X coverage of the estimated genome, from the K-mer distribution using GenomeScope (110 Mb). The final genome assembly spanned 153.7 Mbp (30,095 scaffolds), and the longest contig and N50 length were 41,023 bp and 6,674 bp, respectively (Table 1, Supplementary Table 1). This sequencing was performed after extensive screening for contamination, which is especially critical in this work, since the specimens were collected from the wild. The initial step of the screening was based on BlobTools visualizations ^41^, and scaffolds were removed based on similarity to bacterial sequences with low RNA-Seq coverage (most scaffolds < 10^-1^) and GC contents lower than 40% or higher than 60% (Supplementary Fig. 1). The BUSCO completeness score of the genome assembly was 92.7% (eukaryote dataset), and we observed high mapping ratios for DNA-Seq and RNA-Seq data (Supplementary Table 2). Forty-two megabases (27%) of this genome was inferred to consists of unknown repeats. We further predicted 73 ribosomal genes and 1,081 transfer RNA genes. A total of 42,608 protein-coding genes (maximum TPM > 1) were predicted, of which 57.6% of the genes showed BLASTp homology against the Swiss-Prot database (E-value < 1e-5). Since the assembly length exceeds the expected genome size, this assembly and gene annotations presumably contain duplicate assemblies. Transcriptome assembly was also filtered by CPM (count per million; CPM > 1) to eliminate possible contamination and was further verified by BlobPlot (Supplementary Table 3, Supplementary Fig. 2,). A total of 98.6% of the transcriptome assembly was determined to match the genome assembly by a BLASTn search at an e-value threshold of 1e-50. The genome assembly is not very contiguous, which is a limitation of the ultra-low-input protocol from a single specimen that only contains less than 50 pg of DNA. On the other hand, agreement between the genome and transcriptome assembly, as well as the BUSCO completeness, shows sufficient completeness of the gene repertoire (Supplementary Table 1).

**Table 1.** Statistics of genome assembly.

| Genome Statistics | Before screening<br>(Original) | After screening |
| --- | --- | --- |
| Sequencer | MiSeq |  |
| Scaffold number | 107,804 | 30,095 |
| Total scaffold length (bp) | 388,783,196 | 153,718,311 |
| Average scaffold length (bp) | 3,606 | 5,107 |
| Longest scaffold length (bp) | 326,619 | 41,023 |
| Shortest scaffold length (bp) | 1,000 | 1,001 |
| N50 | 4,891 | 6,674 |
| BUSCO completeness (%) | 89.4 | 87.8 |

**Table 1.** Gene expression and Fold Index of EtAHS proteins. Shown TPM values in Active and Tun are an average of 3 replicates, respectively.

| Gene ID | TPM (average) |  | Fold Index | Folding type |
| --- | --- | --- | --- | --- |
|  | Active | Tun |  |  |
| g97955.t1 | 583 | 400 | -0.210 | EtAHS A |
| g110942.t1 | 504 | 569 | -0.204 |  |
| g158834.t1 | 715 | 825 | -0.215 |  |
| g8031.t1 | 4,146 | 2,538 | -0.092 | EtAHS B |
| g94762.t1 | 4,171 | 2,460 | -0.101 |  |
| g184550.t1 | 3,749 | 2,258 | -0.082 |  |

Next, we compared the *E. testudo* gene set with other sequenced tardigrades: transcriptome data of Heterotardigrada *E*. cf. *sigismundi* and Eutardigrada *Richtersius coronifer*, and genome data of Eutardigrada *R. varieornatus* and *H. exemplaris*. A total of 12,917 genes (1e-25) of *E. testudo* were conserved in all 5 species, and approximately 529 genes were shared only among the heterotardigrades *E. testudo* and *E*. cf. *sigismundi* (Figure 1a). The ratio of conserved genes between each pair of five species showed that the two classes of Tardigrada, Heterotardigrada and Eutardigrada, possess class-specific conservation patterns (Figure 1b), supporting the possibility of distinct anhydrobiosis machineries.

**Figure 1.**
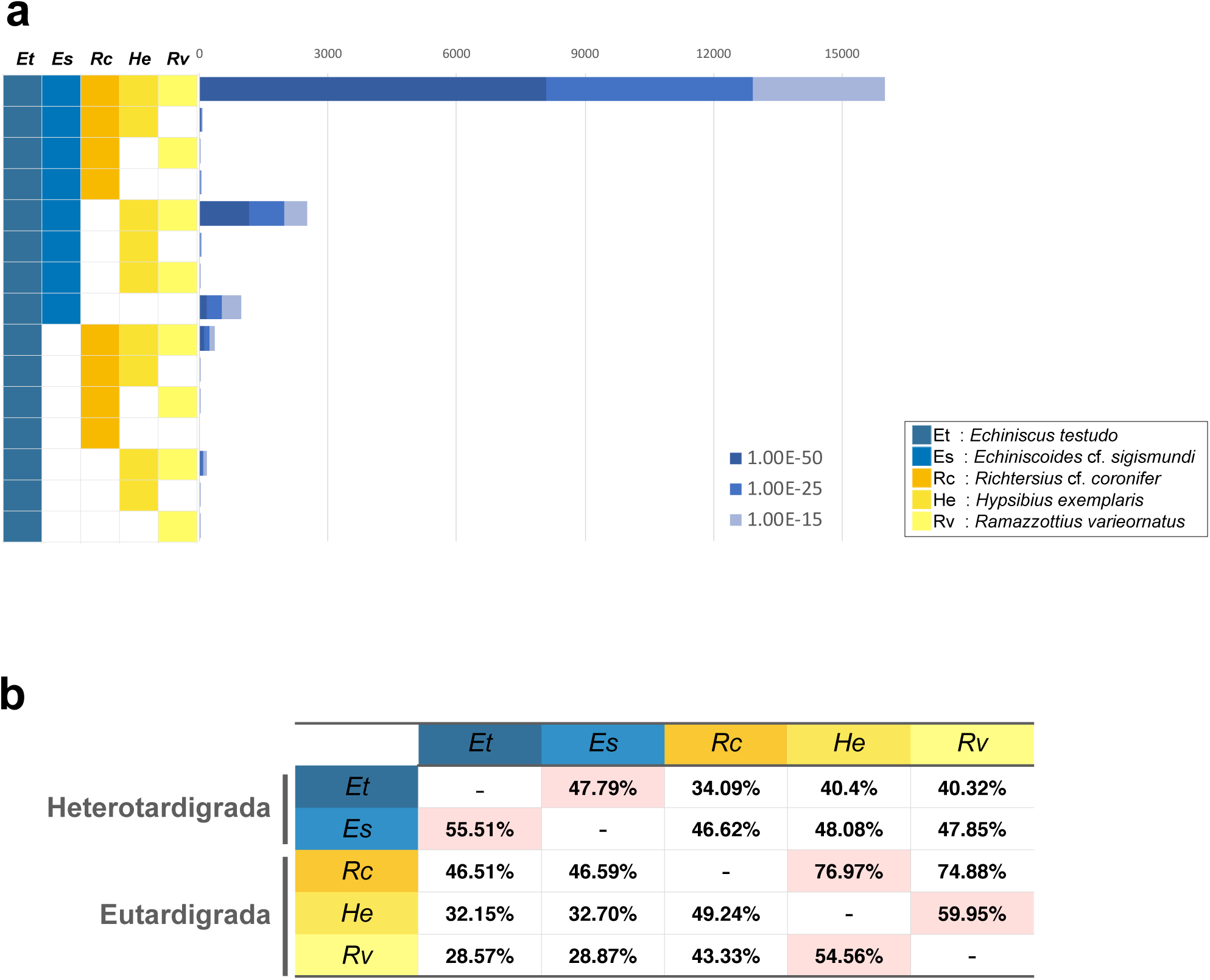
Gene conservation of *Echiniscus testudo* and other tardigrades. Gene distribution of ortholog genes by BLASTp search against Swiss-Prot database. (b) Percentage of gene conservation on E-value < 1E-25. The genomic and transcriptomic data used are shown in the Methods section. Et: *Echiniscus testudo*, Es: *Echiniscoides sigismundi*, Rc: *Richtersius coronifer*, He: *Hypsibius exemplaris*, Rv: *Ramazzottius varieornatus*.

### Comparative genomics of anhydrobiosis in tardigrades

A previous comparative study on the anhydrobiotic machinery between *R. varieornatus* and *H. exemplaris* reported several pathways that contribute to desiccation tolerance ^23^; however, the conservation of these pathways (or genes) has not been identified to date in *Echiniscus*. We first focused on antioxidative stress-related pathways, since previous species have undergone extensive gene loss in H_2_O_2,_ generating stress-responsive pathways, as well as gene duplications in antioxidative stress genes, suggesting intensive genomic adaptations for coping with oxidative stress^23,26^. Naive screening of catalase, superoxide dismutases (SODs), glutamate-S transferase (GST), and heat shock protein (HSP) resulted in 5, 31, 94 and 57 copies, respectively. However, since the genome assembly likely contained duplicated assembly artifacts, we further confirmed these candidates by constructing phylogenetic trees of each gene from genomic and transcriptomic sequences, thereby removing nearly identical orthologs (> 99% identity) and possible artifacts (Supplementary Fig. 3). As a result, we identified 1, 17, 70 and 40 orthologs of catalase, SODs, GST and HSP, respectively (Figure 2). We also observed that several genes related to stress signaling, *TSC2* and *Rheb*, lost in the Eutardigrade linage were conserved in *E. testudo* (Figure 2 and Supplementary Table 4), and this conservation pattern was in line with findings observed for *E*. cf. *sigismundi*, suggesting gene loss in Eutardigrada after divergence from Heterotardigrada. The majority of the genes associated with the stress response (*i*.*e*., loss of Sestrin and HIF-1α) and antioxidative stress genes (*i*.*e*., duplication of SOD and GST) showed conservation patterns in *E. testudo* similar to those observed in *R. varieornatus* and *H. exemplaris*, suggesting a very early trait of the common ancestor of Tardigrada. We subsequently focused on tardigrade-specific anhydrobiosis-related genes that have been identified eutardigrades (*i*.*e*., CAHS, secretory abundant heat soluble (SAHS), mitochondrial abundant heat soluble (MAHS), LEAM and damage suppressor (Dsup)). Similar to *E*. cf. *sigismundi*, these genes were not confirmed in the *E. testudo* genome (BLASTp search, E-value < 1e-3, Figure 2). These results suggest that while there are conserved core mechanisms in the oxidative stress response and stress signaling within the phylum Tardigrada, distinct mechanisms exist in heterotardigrades and eutardigrades.

**Figure 2.**
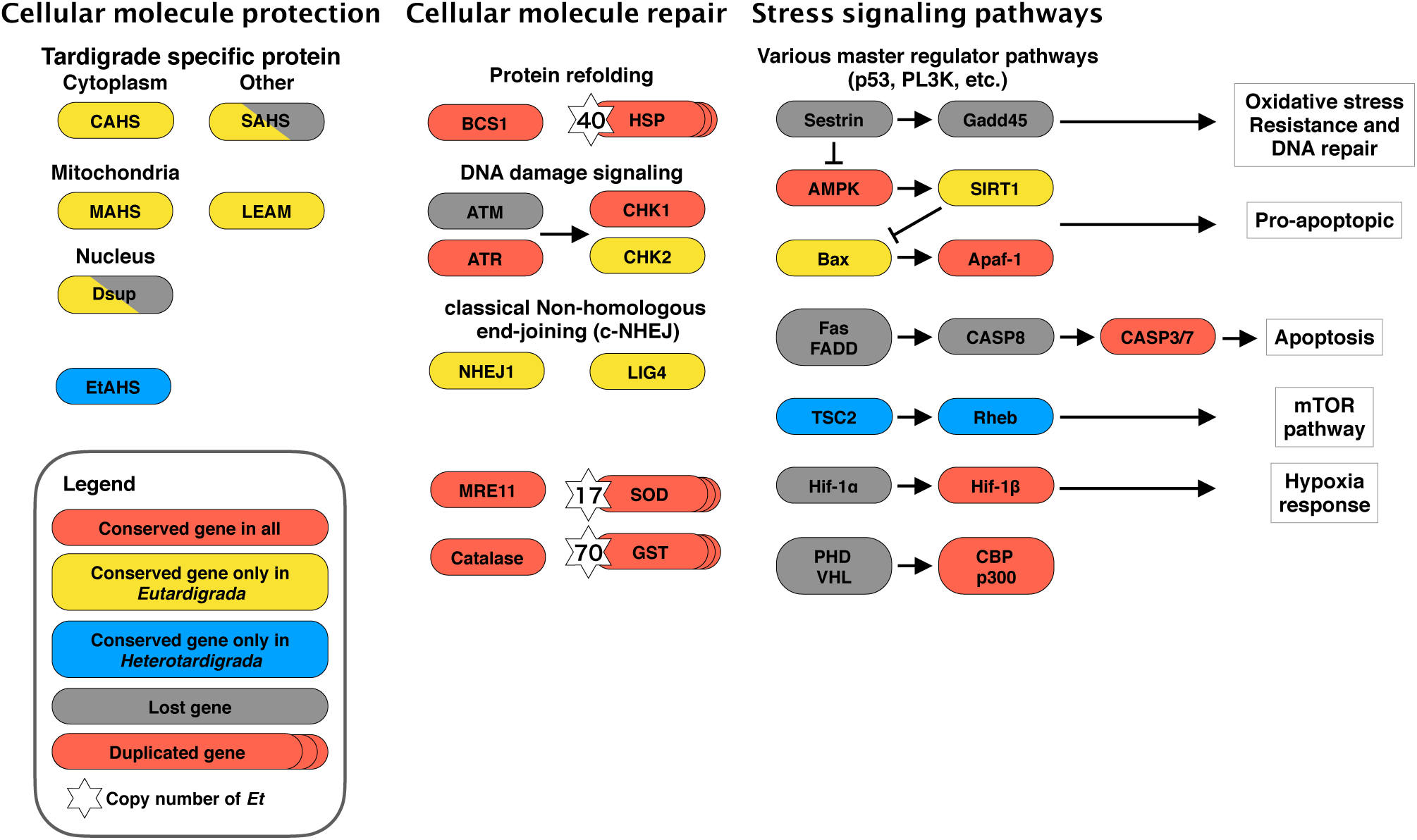
Gene loss and duplication in tardigrades. Coral orange and gray boxes represent genes conserved and lost in all 5 species used in this study. Furthermore, yellow boxes and blue boxes indicate genes conserved only in eutardigrades and in heterotardigrades, respectively. The numbers on the left of the boxes show copy numbers of multiple copy genes in *E. testudo*. AMPK, 5’ adenosine monophosphate-activated protein kinase; Apaf-1, apoptotic protease-activating factor 1; ATM, ataxia-telangiectasia mutated; ATR, ataxia telangiectasia and Rad3-related protein; BCS1, mitochondrial chaperone BCS1; CAHS, cytosolic abundant heat soluble; CASP3/7/8, caspase 3/7/8; CBP, CREB binding protein; CHK1/2, checkpoint kinase 1/2; Dsup, damage suppressor; Fas; tumor necrosis factor receptor superfamily member 6; FADD, Fas-associated death domain; Gadd45, growth arrest and DNA damage-inducible; GST, glutathione S-transferase; Hif1α, Hypoxia-inducible factor 1-alpha; Hif1β, aryl hydrocarbon receptor nuclear translocator; HSP, heat shock protein; LEAM, late embryogenesis abundant protein mitochondrial; MAHS, mitochondrial abundant heat soluble; mTOR, mechanistic target of rapamycin; PHD, plant homeodomain; p300, Histone acetyltransferase; Rheb, GTP-binding protein Rheb; SAHS, secretory abundant heat soluble; SIRT1, sirtuin 1; SOD, superoxide dismutase; TSC1/2, tuberous sclerosis 1/2; VHL, von Hippel-Lindau tumor suppressor.

To screen for heterotardigrade-specific anhydrobiotic machinery, we subsequently analyzed differential gene expression between fully hydrated and desiccated individuals. Differentially expressed genes (DEGs) were extremely limited, with only 21 (q < 0.05) or 13 (q-value < 0.01) candidates being identified (Supplementary Table 5 and Supplementary Fig. 4). The fold change of DEGs was moderately high (median 6.88), but most of these genes had very low expression levels (hydrated TPM < 10 is 16/21 at q < 0.05, 11/13 at q < 0.01), thus resulting in high fold changes. Overall, there is only minimal change in expression between the hydrated and anhydrobiotic states, and screening from differential expression does not seem feasible, as is observed for the strong anhydrobiote *R. varieornatus* ^26^.

### Novel heat soluble protein in heterotardigrades

The results obtained to date suggest the presence of heterotardigrade-specific machinery of anhydrobiosis, which complements the functions of the LEA or CAHS proteins observed in other anhydrobiotes, presumably contributing to cellular molecule protection upon desiccation ^27^. Therefore, we considered heat solubility screening to obtain analogous proteins. Following heat-soluble proteomics (Supplementary Table 6), filtering for highly expressed genes (maximum TPM > 100) from transcriptome data and lack of sequence similarity in Swiss-Prot yielded 32 genes as final candidates (Supplementary Table 7). These candidates were further screened for their disordered nature with negative fold index values, as these genes are predicted to have large intrinsically unstructured regions. This approach resulted in six genes of two fold index patterns, containing a high proportion of predicted alpha helical regions, similar to those of CAHS1 (Table 2, Figure 3). Although there is no sequence similarity, these *Echiniscus*-specific, heat-soluble, constitutively highly expressed (these genes were not DEGs in the above analysis and are highly expressed both in hydrated and tun states), alpha-helical proteins with intrinsically unstructured characteristics are all analogous to the CAHS protein identified in *R. varieornatus*; therefore, we named these genes EtAHS (*Echiniscus testudo* Abundant Heat Soluble). A BLAST search also identified EtAHS in *E*. cf. *sigismundi*, but it was not found in all eutardigrades, that is, *R. varieornatus, H. exemplaris* and *R. coronifer*, suggesting heterotardigrade-specific conservation.

**Figure 3.**
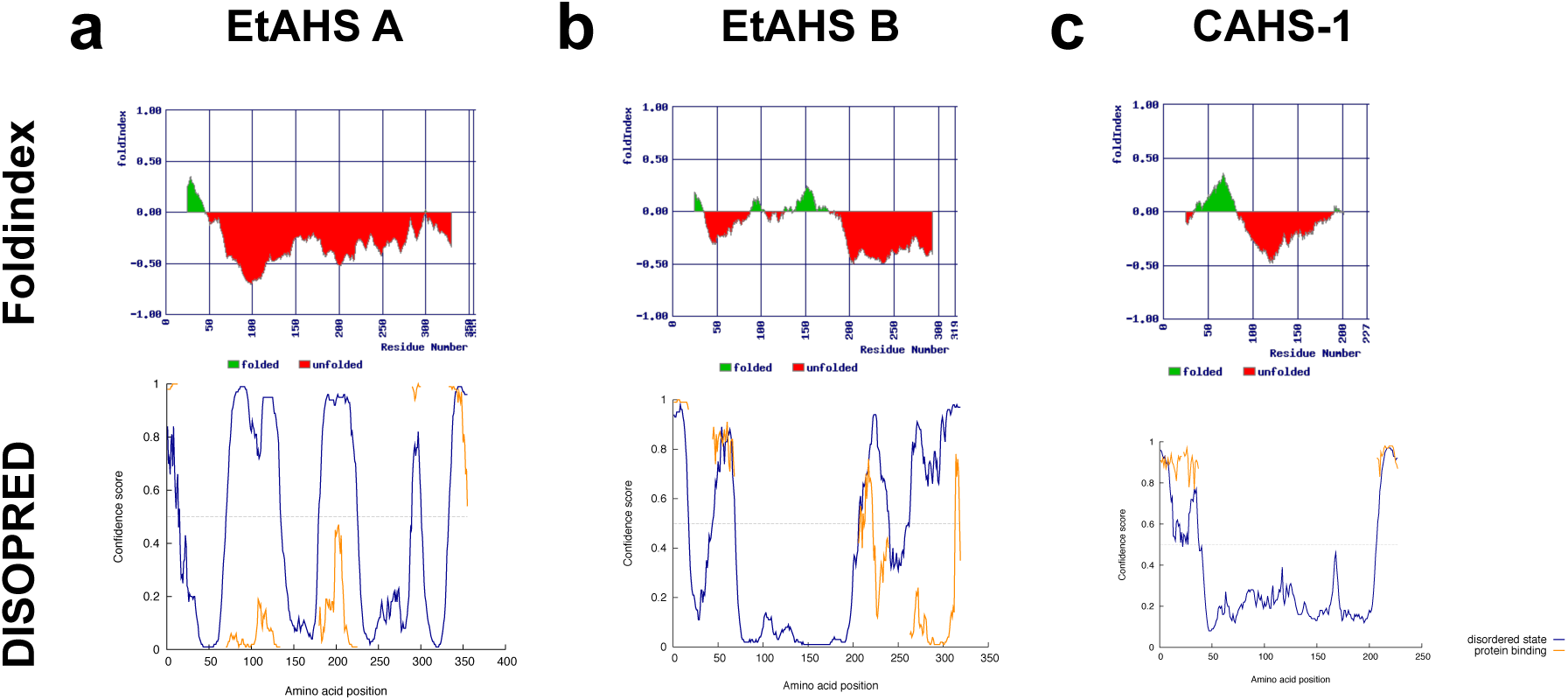
Folding prediction of EtAHS proteins. Folding of (a) EtAHS A, (b) EtAHS B and (c) CAHS1 proteins. The upper tier shows Fold Index and the lower tier shows DISOPRED folding. In Foldindex, positive (green area) and negative (red area) numbers represent ordered and disordered protein, respectively. Amino acids suggested as being folded are shown in the green area and unfolded in the red area. In DISOPREAD, the blue line indicates disordered state, and the orange line indicates protein binding.

### Structural analysis of EtAHS proteins

We next performed structural characterization of the recombinant EtAHS proteins using NMR and CD spectroscopy (Figure 4). The ^1^H-^15^N HSQC peaks were observed within a narrow spectral region (7.4-8.4 ppm for ^1^H chemical shift), indicating that the EtAHS proteins were largely unstructured in solution. The unstructured properties of EtAHS proteins were confirmed by CD spectral data. Moreover, the CD data demonstrated that the desolvating agent of trifluoroethanol (TFE) induced alpha-helical conformation in the EtAHS proteins. These properties are similar to those of CAHS, which also undergoes a conformational transition from a largely unstructured state to an alpha-helical state in water-deficient conditions.

**Figure 4.**
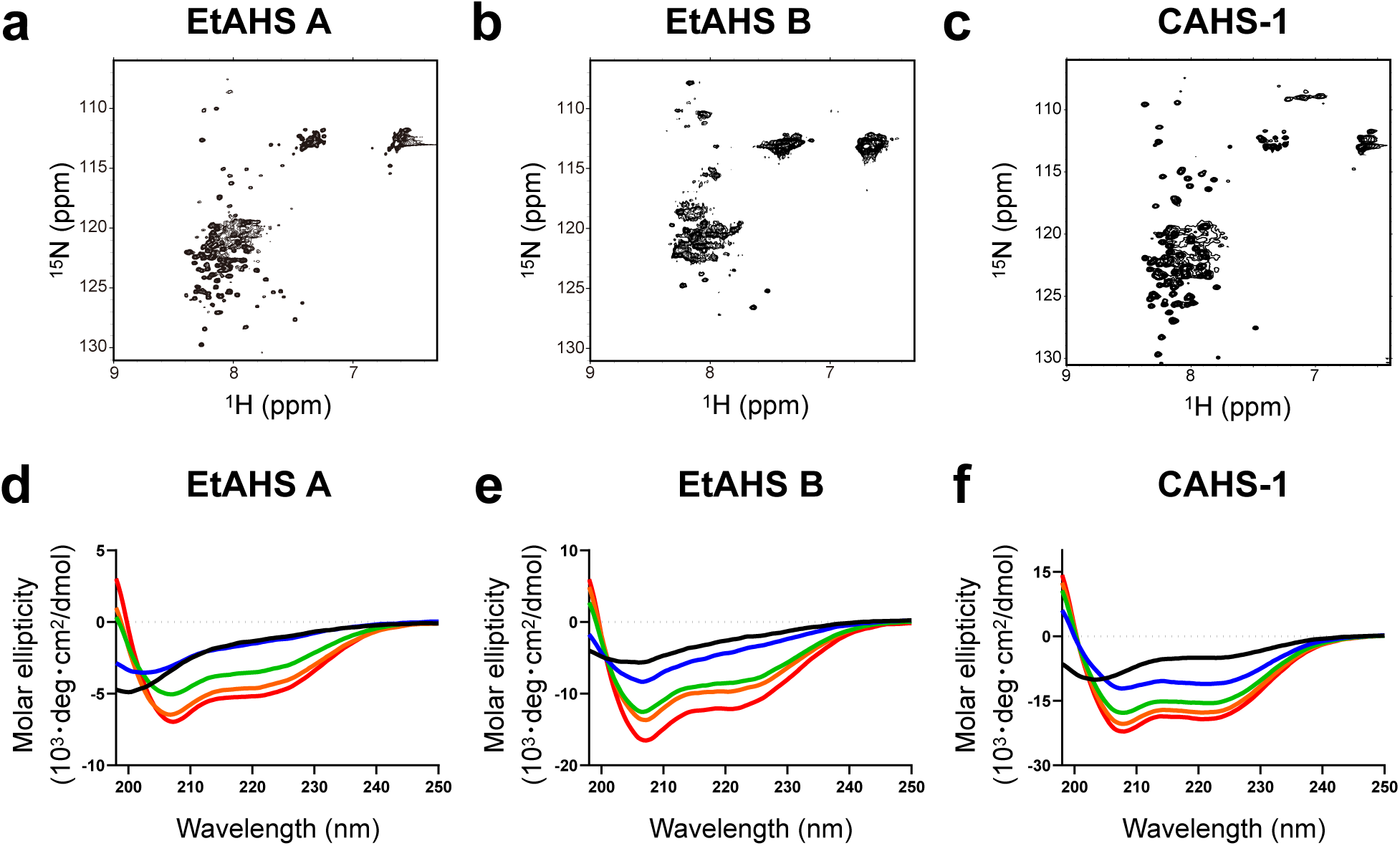
Structural analysis of EtAHS proteins. ^1^H-^15^N HSQC spectra of (a) EtAHS A, (b) EtAHS B, and (c) CAHS1 proteins. CD spectra of (d) EtAHS A, (e) EtAHS B, and (f) CAHS1 proteins in the absence (black) and presence of 10% (blue), 30% (green), 50% (orange), and 70% (red) (v/v) TFE.

## Discussion

Molecular and genomic studies of tardigrades have mostly been limited to the class Eutardigrada, where numerous species are culturable in laboratory conditions. On the other hand, no sustainable culture of heterotardigrades has been established to date. Therefore, we employed an ultra-low-input genome sequencing protocol that we developed previously ^38-40^ to achieve whole genome sequencing from a single specimen of tardigrade containing less than 50 pg of DNA collected from the wild. However, such a sample requires strong precautions to minimize the possibility of contamination ^23,38,42^. After rigorous filtering of possible contaminants as described above, we generated a 153.7 Mbp draft genome assembly of *E. testudo* with high genome completeness (BUSCO score; 92.7%) and a 98.6% match of transcriptome assembly to the genome. This report is the first to describe a heterotardigrade genome, and this novel resource may facilitate future comparative genomics and evolutionary studies of the phylum Tardigrada. Transcriptome assembly of a heterotardigrade, *E*. cf. *sigismundi*, has been reported ^25^, and while such data are valuable in identifying the presence of a gene, genomic data are required to discuss the absence of a gene and to perform further analysis of other noncoding features.

Comparing the gene repertoire of *E. testudo* with other tardigrades, namely, *E*. cf. *sigismundi, R. varieornatus, H. exemplaris* and *R. coronifer*, duplication of stress-response genes, such as GST, SODs and HSP, was observed as a common characteristic of anhydrobiotic tardigrades. Reactive oxygen species and the oxidative damage induced by them are among the most critical sources of cellular damage upon desiccation; therefore, the expansion of the antioxidant defense repertoire appears to be a common mechanism in anhydrobiosis, not one that is limited to tardigrades ^43,44^. In addition, we identified several genes in stress signaling pathways, i.e., genes involved in stress signaling pathways. Sestrin, Gadd45, FADD, and Caspase 8 were lost *E. testudo*, similar to the eutardigrades *R. varieornatus* and *H. exemplaris*, suggesting that these orthologs were lost prior to the diversion of Eutardigrada and Heterotardigrada. Hypoxia-inducible factors (HIFs) are transcription factors that serve as master regulators of oxygen homeostasis and are highly conserved in most metazoans, including desiccation tolerance species ^45^. Initially, the loss of this modulator of apoptosis induction was suggested to be associated with anhydrobiosis to prevent apoptosis and to enable cellular repair upon rehydration from anhydrobiosis ^26^. However, if the loss of this gene occurred before the divergence of Heterotardigrada and Eutardigrada, this would imply that this gene loss is also present in the marine tardigrades of Arthrotardigrada, which do not possess anhydrobiotic capability, thereby weakening the link between these gene losses and anhydrobiosis. Recently, it was reported that an abundant intertidal crustacean, the copepod *Tigriopus californicus*, also lacks HIF-α and EGLN, but this species can tolerate extremely low levels of available O_2_ for at least 24 h ^46^. This finding suggests that the loss of HIF-α/EGLN is not specific to tardigrades and may not be related to anhydrobiosis.

While these core components are conserved, none of the tardigrade-specific proteins related to anhydrobiosis identified in *R. varieornatus*, namely, CAHS, SAHS, MAHS, LEAM, and Dsup, is observed in *E. testudo*. CAHS protein is highly expressed in *R. varieornatus*, and the presence of a large number of paralogs within the genome suggests an important role played by this protein during anhydrobiosis. While CAHS is constitutively highly expressed in the strong anhydrobiote *R. varieornatus*, expression of this protein is induced upon desiccation in the weak anhydrobiote *H. exemplaris*, indicating a direct functional link in anhydrobiosis ^23^. Therefore, analogous function is expected in *E. testudo* but without sequence similarity or homology, as in the case of different LEA groups ^47^. The rate of gene conservation is higher within the classes of Tardigrada, and 529 genes are only shared by heterotardigrades. Eighty-nine of these proteins have a homolog in Swiss-Prot (1e-25); therefore, candidates for heterotardigrade-specific anhydrobiosis-related proteins should be among these 440 genes. Screening of these candidates by differential expression in active and tun states was not possible due to the lack of overall expression change. This result is not surprising because strong anhydrobiotes, such as *R. varieornatus* and *E. testudo*, can enter anhydrobiosis immediately, forming tuns and losing most of their body water within 30 min. This time interval is not sufficient to conduct de novo gene transcription and protein translation, and a lack of expression changes has been reported for *R. varieornatus*, where anhydrobiosis-related proteins are constitutively highly expressed ^23^.

Therefore, we performed heat solubility proteomics assays to identify hydrophilic proteins, following the procedures established for the identification of LEA and CAHS proteins ^19,27^. These classes of proteins are notably hydrophilic and are often comprised of intrinsically unstructured domains that turn into alpha-helical structures upon water loss. These properties enable the protein to remain soluble even after heat treatment at near-boiling temperatures. As a result, we identified two families of six genes that we designated EtAHS that were conserved only in heterotardigrades. These proteins were predicted to contain unstructured regions similar to those observed in CAHS and SAHS, suggesting that these genes may be analogous ^27^. Further confirmation of the EtAHS structure using recombinant proteins expressed in *E. coli* showed a disordered nature by NMR spectroscopy and a change in structure from random coil to alpha helix upon water loss, which was emulated by replacing water with TFE, and both structural features resembled those of CAHS. The abundance, heat solubility, hydrophilic nature, and intrinsically disordered structure that forms an alpha-helical structure upon water loss are all characteristic of the LEA protein family. LEA proteins do not exhibit sequence similarity among groups ^47,48^; therefore, it is possible to consider that tardigrade heat-soluble protein families may be a new LEA protein family group, but such generalization is beyond the scope of this work and is a possible direction for future research. Moreover, the role played by EtAHS or CAHS in anhydrobiosis has not been elucidated to date. Few studies have investigated this topic, and they have obtained ambiguous results ^49^. Further studies on this subject are also necessary to fully elucidate the molecular mechanisms underlying anhydrobiosis in tardigrades. Nevertheless, the existence of analogous proteins without any sequence similarity suggests the importance of this class of proteins in anhydrobiosis and that tardigrades have independently evolved these protein families at least two times, once in Heterotardigrada and once in Eutardigrada, to enable anhydrobiosis. This phenomenon may represent an example of convergent evolution and highlights the complex evolutionary history of anhydrobiosis within the phylum Tardigrada.

## Conclusions

In this study, we performed genomic and transcriptomic sequencing of the heterotardigrade *E. testudo* by using an ultra-low-input library sequencing method, and by combining these results with heat-soluble proteome data, we identified a novel heterotardigrade-specific anhydrobiosis-related gene family designated EtAHS, which shares multiple analogous characteristics, including heat solubility, constitutive high expression, intrinsically unstructured nature, and shift to alpha-helical structure upon water loss, with the eutardigrade analog CAHS. Structural features were further confirmed using NMR and CD analyses, which also indicated similar characteristics to those of CAHS. These results suggest the existence of two distinct, independently acquired mechanisms of anhydrobiosis in the two classes of Tardigrada, which may represent an example of convergent evolution.

## Materials and Methods

### Animals

We collected *E. testudo* from moss collected from Otsuka-machi, Tsuruoka city, Japan (38°44′23″N, 139°48′28″E). The moss was soaked in water for more than 3 h, and *E. testudo* specimens were picked and collected on an agar-coated plate. *E. testudo* was allowed to walk around the agar plate for two days, with rigorous washing and renewed plate each day to remove surface and gut contaminants. Tun (anhydrobiotic) state was induced by suspending 20-21 animals in 100 µl of distilled water and by placing it on a 2-cm x 2-cm mesh filter on a paper towel (ASONE). The animals were desiccated at 35% RH for 24 h controlled by 85% glycerol solution ^50^, followed by 0% RH at room temperature for complete desiccation. Anhydrobiotic success was determined by an over 90% recovery rate after 24 h.

### Genome and transcriptome sequencing

We used the ultra-low-input library sequencing protocol established in our previous study ^39,40^. In brief, genomic DNA was extracted using a Quick-gDNA Microprep Kit (Zymo Research) after lysis with two freeze-thaw cycles of −80°C and 37°C incubation, and the extracted DNA was sheared to 550-bp target fragments with Covaris M220. The Illumina sequencing library was constructed with a Thruplex DNA-Seq kit (Rubicon Genomics), and the purified library was quantified using a Qubit Fluorometer (Life Technologies). The sequencing library was sequenced on the MiSeq platform (Illumina) with 300 bp paired settings. For RNA-Seq, the total RNA was extracted from active and tun samples (3 replicates) using Direct-Zol RNA MicroPrep Kits (Zymo Research), and the sequencing library was constructed using SMARTer v4 Ultra Low Input RNA Kit for Sequencing (Clonetech) and KAPA HyperPlus Library Preparation Kit (KAPA BIOSYSTEMS). The RNA-Seq library was multiplexed and sequenced on the NextSeq 500 platform (Illumina) 300 cycles High Output Mode (paired-end). De-multiplexing of RNA-Seq reads was conducted using bcl2fastq v.2 (Illumina). Both DNA-Seq and RNA-Seq reads were submitted to FastQC ^51^ for quality validation.

### Genome assembly and gene finding

Following genome size estimation by GenomeScope ^52^, the MiSeq reads were assembled with MaSuRCA (v3.1.3) ^53^ with the following parameters (PE 350 150). Since the specimens were directly collected from the wild, we needed to be extra careful about possible contamination in the sequences. To validate contamination in the genome assembly ^23,38,42^, we first mapped the MiSeq DNA-Seq reads to the assembled genome with BWA (v0.7.17) ^54^ and converted the output files with SAMtools (v1.7) ^55^. Next, we conducted a BLASTn search against the NCBI nr database ^56,57^. These results were submitted to BlobTools (v1.0) ^41^ for visualization. Contigs with GC content below 40% and above 60% and contigs identified to be bacterial were determined as contaminants and removed from the assembly. We validated the genome completeness with BUSCO (v3.0.2) using the eukaryote lineage gene set ^58^.

For identifying genes, we first mapped the RNA-Seq data (active and tuned three replicates each) to the genome assembly with TopHat2 (v2.1.1) ^59^. The resulting BAM files were submitted to BRAKER (v1.9) with default settings ^60^. To remove possible bacterial contamination, we validated whether these query transcripts were included in the transcriptome assembly produced with Trinity v.2.5.1 ^61^ since polyA selection during the sequencing library construction process would filter non-polyA transcripts. Furthermore, we performed tBLASTn against the nr database and excluded nonmetazoan hits to remove possible eukaryotic contamination. We submitted these query sequences to a tBLASTn search against the transcriptome assembly and excluded transcripts that had identity below 99%. To annotate the predicted gene models, we performed similarity searches using BLASTp (E-value < 1e-5) against Swiss-Prot ^62^ and against tardigrade-predicted proteome sequences of *R. varieornatus* and *H. exemplaris* genomes (E-value < 1e-5). We also submitted the amino acid sequences to KEGG Automatic Annotation Server ^63,64^ to assign KEGG orthology IDs. De novo repeat region identification was conducted by RepeatScout ^65^ and RepeatMasker ^66^. We searched tRNA using tRNAscan ^67^and rRNA using RNAmmer ^68^.

This final assembly data contained 42,608 predicted protein-coding genes with expression levels of > 1 transcript per million (TPM) in at least one of six RNA-Seq samples. A total of 57.6% of the genes showed BLASTp similarity against the Swiss-Prot database (E-value < 1e-5). To identify genes that were induced during anhydrobiosis, we quantified the gene expression (TPM) with Kallisto (v0.42.1) ^69^. Additionally, we mapped the RNA-Seq reads to the coding sequences with BWA MEM and tested for differential expression using DESeq2 ^54,70^. Transcripts with FDR below 0.05 were defined as DEGs.

### Gene expansion and lost pathway analysis

To validate anhydrobiosis-related genes reported in previous tardigrade genomic studies ^23,26^, KAAS orthologous annotation was used to initially screen for possible gene losses. For orthologs that were initially found to be missing in the *E. testudo* genome, we further confirmed gene loss by using orthologs in *R. varieornatus, H. exemplaris*, and Swiss-Prot for tBLASTn searches to the *E. testudo* genome.

To identify multicopy genes, we conducted multiple alignment and counted the copy number. Multiple alignment with *R. varieornatus* and *C. elegans* was carried out using MAFFT (v7.221) ^71,72^. Phylogenetic trees were constructed using FastTree (v2.1.10) with default options ^73^ and visualized with Interactive Tree of Life (iTOL) ^74^. Regarding extremely similar sequences, genes with BLASTn identity > 99% were regarded as identical artifact copies due to possible misassembly (Supplementary Table 5, Supplementary Fig. 1 phylogenetic tree).

### Heat soluble proteomics

Heat soluble proteomics was conducted as previously described ^27^. Briefly, approximately 3,000 individuals were collected from the wild and cleaned as described above and homogenized using BioMasher II (Nippi) in PBS (Nippon Gene) on ice with protease inhibitors (Roche). The lysate was heated at 92°C for 20 min, and the soluble fraction was collected by taking the supernatant after centrifugation at 12,000 rpm for 20 min. Proteins were digested with trypsin, and tryptic peptides were separated and identified with an UltiMate 3000 nanoLC pump (Dionex Co., Sunnyvale, CA, USA) and an LTQ Orbitrap XL ETD (Thermo Electron, Waltham, MA, USA). Corresponding peptide sequences were retrieved from six frame translation data of our initial genome assembly using MASCOT software (Matrix Science) ^75^. Candidates were further screened with the following conditions: (1) lack of conservation in other tardigrades or metazoans and (2) high mRNA expression (TPM >100) in the tun state. We then predicted the structural features using the Fold Index ^76^ and DISOPRED ^77^.

### Structural analysis of EtAHS proteins

The gene derived from g97955.t1, which encodes residues Thr29-Gln355 of EtAHS A protein, was cloned into pET28a (Novagen). The gene derived from g8031.t1, which encodes residues Gln20-Lys319 of EtAHS B protein, was cloned into modified pCold-1 (Takara Bio Inc.), in which the factor Xa cleavage site was replaced with tobacco etch virus (TEV) protease cleavage. The recombinant EtAHS proteins were expressed in the *E. coli* BL21(DE3) CodonPlus strain cultured in LB medium. The His_6_-tagged fusion EtAHS A protein was purified from cell lysates with a Ni^2+^-nitrilotriacetic acid column (GE Healthcare). The fusion protein was cleaved by incubation with thrombin protease (Sigma Aldrich) and then purified with Superdex 200 pg (GE Healthcare). For preparation of EtAHS B protein, after lysis by sonication, the insoluble inclusion bodies were extensively washed with 20 mM Tris-HCl (pH 8.0) containing 150 mM NaCl and 2% Triton X-100 and then solubilized with 6 M guanidinium chloride, 50 mM Tris-HCl (pH 8.0), and 1 mM dithiothreitol. The solubilized proteins were refolded by dilution (0.1 mg/mL) in 20 mM Tris-HCl (pH 7.5), 100 mM NaCl, 3 mM reduced glutathione, and 0.3 mM oxidized glutathione at 4°C for 12 h. The His_6_-tagged fusion protein was purified with a Ni^2+^-nitrilotriacetic acid column (GE Healthcare). The EtHAS B protein, from which the N-terminal His_6_-tag peptide was removed by TEV protease digestion, was further purified with a HiLoad Superdex 75 pg (GE Healthcare). The expression and purification of recombinant CAHS1 protein (*R. varieornatus*, UniProt ID: J7M799) were performed as described in the literature with slight modifications ^27^. For the production of ^15^N-labeled EtAHS A, EtAHS B, and CAHS1 proteins, cells were grown in M9 minimal media containing [^15^N]NH_4_Cl (1 g/L). CD spectra were measured at room temperature on a JASCO J-720WI apparatus using a 1.0-mm path length quartz cell. The EtAHS A, EtAHS B, and CAHS1 proteins were dissolved at protein concentrations of 5.6 µM, 2.3 µM, and 6.8 µM, respectively, in 10 mM potassium phosphate buffer (pH 7.4) in the absence and presence of TFE.

NMR spectral measurements were made on a Bruker DMX-500 spectrometer equipped with a cryogenic probe. The probe temperature was set to 5°C. ^15^N-labeled EtAHS A, EtAHS B, and CAHS1 proteins were dissolved at a protein concentration of 0.15 mM in 10 mM potassium phosphate buffer (pH 7.4) containing 5% (v/v) ^2^H_2_O.

## Data availability

The genome/transcriptome sequencing data sets are available at the BioProject accession number PRJNA669587. The assembly and coding sequencing data are available at https://doi.org/10.6084/m9.figshare.13060634.

## Acknowledgments

The authors are grateful to Yuki Takai (IAB) and Naoko Ishii (IAB) for providing experimental support and to Konosuke M. Ii (IAB) for collecting *E. testudo* on proteome analysis. The authors also thank Yudai Sasaki (NCU) for his contribution during the early stage of this study, Dr Tadashi Satoh, Kumiko Hattori (NCU) and Dr Ean Wai Goh (ExCELLS) for spectroscopic measurements. This research was supported by Grants-in-Aid for Scientific Research KAKENHI of the Japan Society for the Promotion of Science (No. 17H03620), the Nanotechnology Platform Program (Molecule and Material Synthesis, JPMX09S19MS0051) of MEXT, Japan, the Joint Research of the Exploratory Research Center on Life and Living Systems (ExCELLS program No.18-207, 19-208, 19-501), and research funds from the Yamagata prefectural government and Tsuruoka city.

## Contributions

K.A. conceived and designed the project. Y.M. conducted the RNA sequencing, data analysis and bioinformatics analysis. M.Y.U. and K.K. performed experiments of NMR and CD spectrum. M.F. conducted proteome analysis. K.A performed genome sequencing. The study was supervised by K.A. and M.T. The manuscript was drafted by Y.M, M.Y.U and K.A, and all authors reviewed and approved the final manuscripts.

