## Supplementary for "Multiomics study of a heterotardigrade, *Echinisicus testudo*, suggests convergent evolution of anhydrobiosis-related proteins in Tardigrada"

### Supplementary Figures

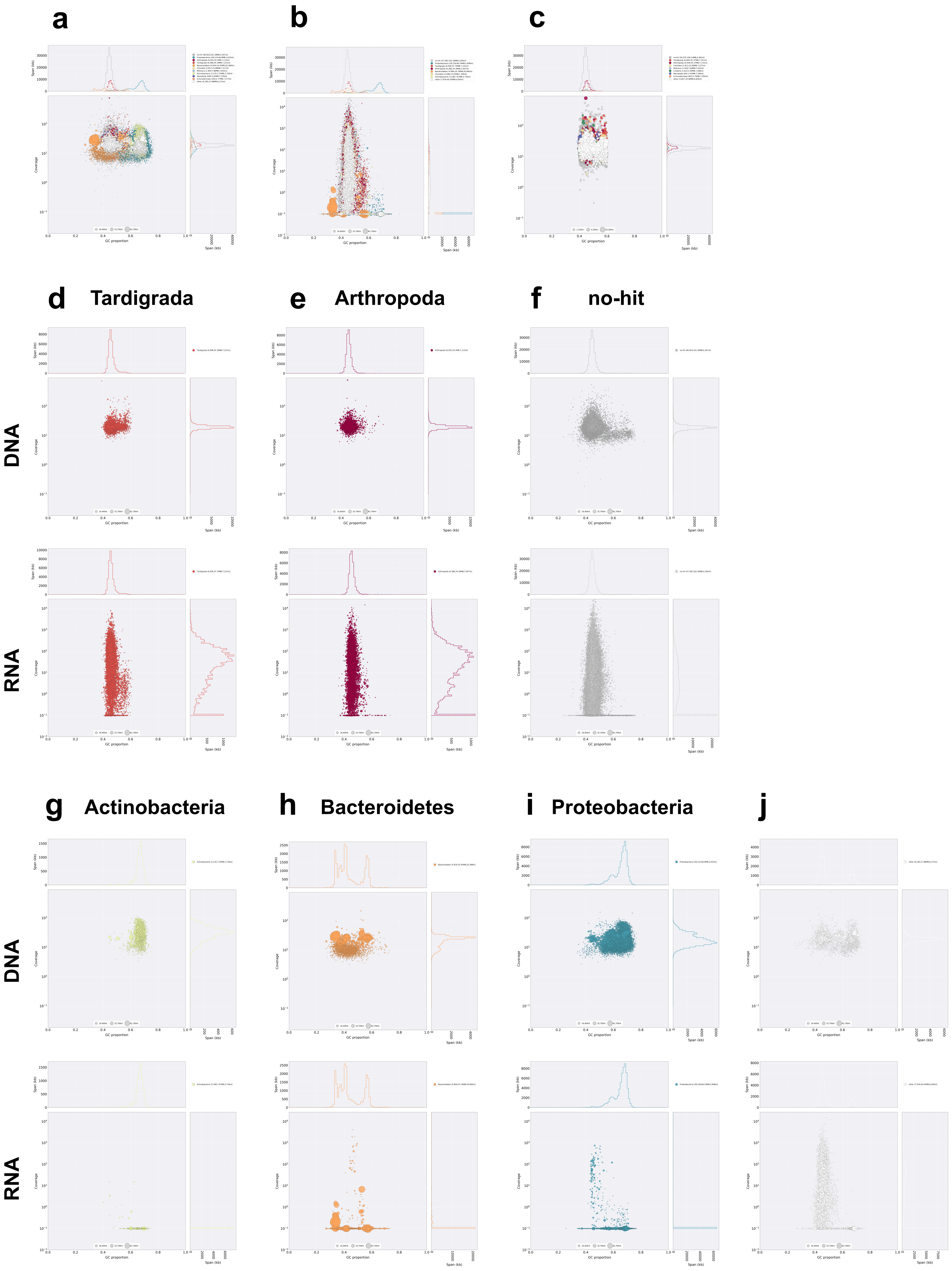

**Supplementary Figure 1. Blobplot analysis of the *E. testudo* genome assembly.**

The assembled genome was subjected to BlobTools for possible contamination identification and visualize as blopblot. (a) Blobplot of original genome assembly (before screening), (b) blobplot of original genome assembly with RNA-Seq coverage data, and (c) blobplot of after screening genome assembly. (d-j) Blobplot printed for each taxonomic group separately. The upper tier was using DNA-Seq data for mapping and the lower tier blobplot was using RNA-Seq data for mapping. (d) Taxonomic group; Tardigrada, (e) Arthropod, (f) "no-hit", (g) Bacteoidetes, (h) Proteobacteria, (i) Actinobacteria and (j) other. Scaffolds were submitted to DIAMOND BLASTX analysis against UniProt Reference Proteomes (2018_09 version) for taxonomy identification, and mapped data by BWA were used for coverage calculation. This information was analyzed by BlobTools.

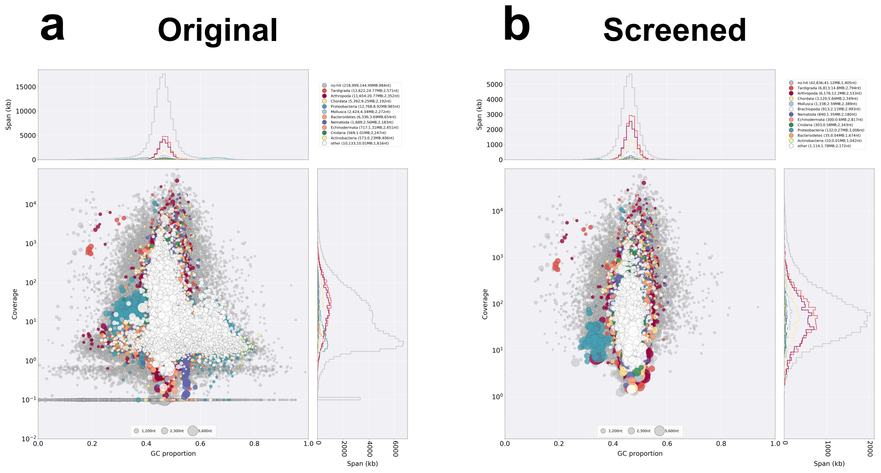


**Supplementary Figure 2. Blobplot analysis of the *E. testudo* transcriptome assembly.**

The assembled transcriptome was subjected to BlobTools for possible contamination identification and visualize as blobplot. (a) Blobplot of original transcriptome assembly data and (b) after screening transcriptome assembly data. Scaffolds were submitted to DIAMOND BLASTX analysis against UniProt Reference Proteomes (2018_09 version) for taxonomy identification, and mapped data by BWA were used for coverage calculation. This information was analyzed by BlobTools.

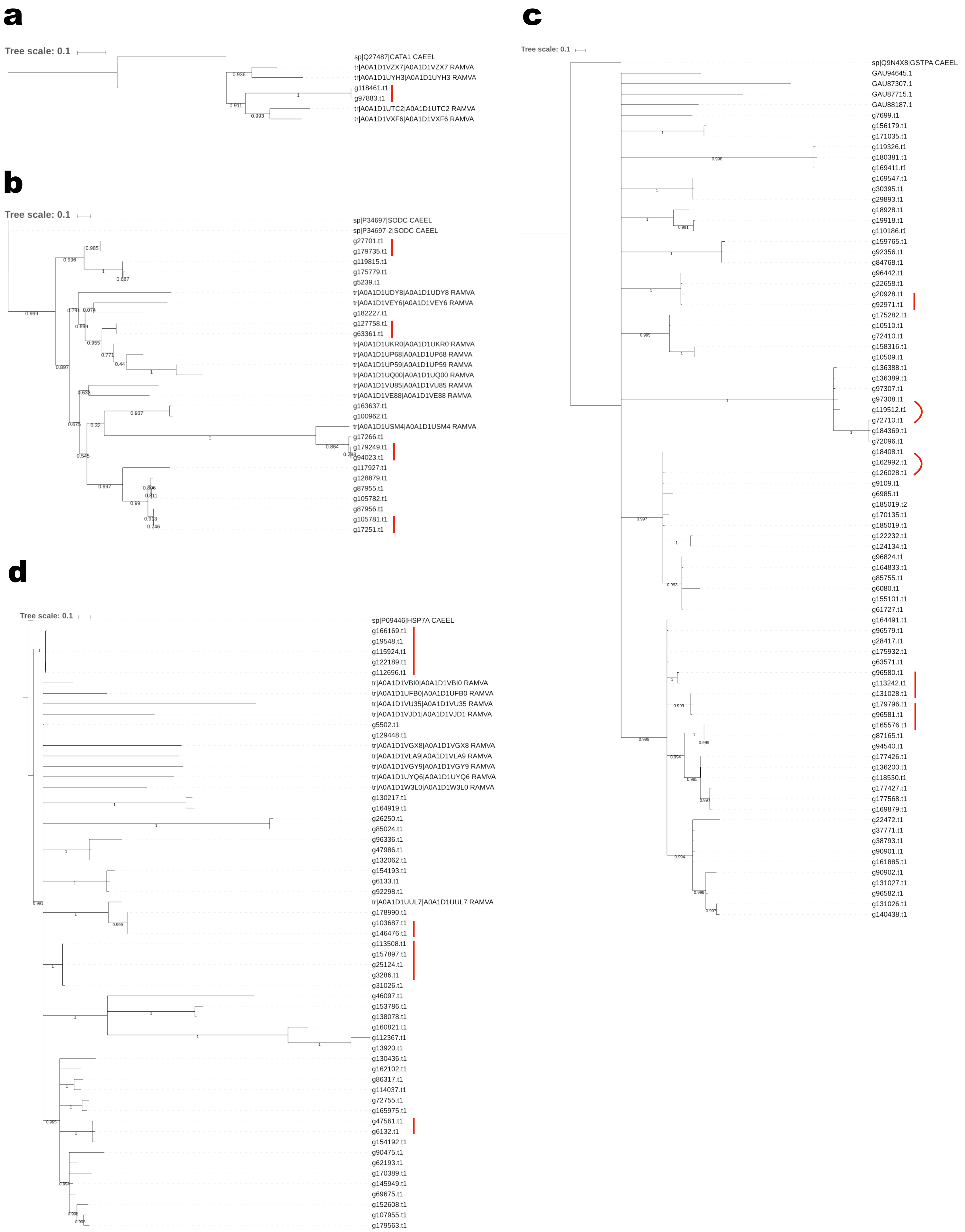
a
Tree scale: 0.1
sp|Q27487|CATA1 CAEEL
tr|A0A1D1VZX7|A0A1D1VZX7 RAMVA
tr|A0A1D1UYH3|A0A1D1UYH3 RAMVA
g118461.t1
g97883.t1
tr|A0A1D1UTC2|A0A1D1UTC2 RAMVA
tr|A0A1D1VXF6|A0A1D1VXF6 RAMVA
b
Tree scale: 0.1
sp|P34697|SODC CAEEL
sp|P34697-2|SODC CAEEL
g27701.t1
g179735.t1
g119815.t1
g175779.t1
g5239.t1
tr|A0A1D1UDY8|A0A1D1UDY8 RAMVA
tr|A0A1D1VEY6|A0A1D1VEY6 RAMVA
g182227.t1
g127758.t1
g63361.t1
tr|A0A1D1UKR0|A0A1D1UKR0 RAMVA
tr|A0A1D1UP68|A0A1D1UP68 RAMVA
tr|A0A1D1UP59|A0A1D1UP59 RAMVA
tr|A0A1D1UQ00|A0A1D1UQ00 RAMVA
tr|A0A1D1VU85|A0A1D1VU85 RAMVA
tr|A0A1D1VE88|A0A1D1VE88 RAMVA
g163637.t1
g100962.t1
tr|A0A1D1USM4|A0A1D1USM4 RAMVA
g17266.t1
g179249.t1
g94023.t1
g117927.t1
g128879.t1
g87955.t1
g105782.t1
g87956.t1
g105781.t1
g17251.t1
c
Tree scale: 0.1
sp|Q9N4X8|GSTPA CAEEL
GAU94645.1
GAU87307.1
GAU87715.1
GAU88187.1
g7699.t1
g156179.t1
g171035.t1
g119326.t1
g180381.t1
g169411.t1
g169547.t1
g30395.t1
g29893.t1
g18928.t1
g19918.t1
g110186.t1
g159765.t1
g92356.t1
g84768.t1
g96442.t1
g22658.t1
g20928.t1
g92971.t1
g175282.t1
g10510.t1
g72410.t1
g158316.t1
g10509.t1
g136388.t1
g136389.t1
g97307.t1
g97308.t1
g119512.t1
g72710.t1
g184369.t1
g72096.t1
g18408.t1
g162992.t1
g126028.t1
g9109.t1
g6985.t1
g185019.t2
g170135.t1
g185019.t1
g122232.t1
g124134.t1
g96824.t1
g164833.t1
g85755.t1
g6080.t1
g155101.t1
g61727.t1
g164491.t1
g96579.t1
g28417.t1
g175932.t1
g63571.t1
g96580.t1
g113242.t1
g131028.t1
g179796.t1
g96581.t1
g165576.t1
g87165.t1
g94540.t1
g177426.t1
g136200.t1
g118530.t1
g177427.t1
g177568.t1
g169879.t1
g22472.t1
g37771.t1
g38793.t1
g90901.t1
g161885.t1
g90902.t1
g131027.t1
g96582.t1
g131026.t1
g140438.t1
d
Tree scale: 0.1
sp|P09446|HSP7A CAEEL
g166169.t1
g19548.t1
g115924.t1
g122189.t1
g112696.t1
tr|A0A1D1VBI0|A0A1D1VBI0 RAMVA
tr|A0A1D1UFB0|A0A1D1UFB0 RAMVA
tr|A0A1D1VU35|A0A1D1VU35 RAMVA
tr|A0A1D1VJD1|A0A1D1VJD1 RAMVA
g5502.t1
g129448.t1
tr|A0A1D1VGX8|A0A1D1VGX8 RAMVA
tr|A0A1D1VLA9|A0A1D1VLA9 RAMVA
tr|A0A1D1VGY9|A0A1D1VGY9 RAMVA
tr|A0A1D1UYQ6|A0A1D1UYQ6 RAMVA
tr|A0A1D1W3L0|A0A1D1W3L0 RAMVA
g130217.t1
g164919.t1
g26250.t1
g85024.t1
g96336.t1
g47986.t1
g132062.t1
g154193.t1
g6133.t1
g92298.t1
tr|A0A1D1UUL7|A0A1D1UUL7 RAMVA
g178990.t1
g103687.t1
g146476.t1
g113508.t1
g157897.t1
g25124.t1
g3286.t1
g31026.t1
g46097.t1
g153786.t1
g138078.t1
g160821.t1
g112367.t1
g13920.t1
g130436.t1
g162102.t1
g86317.t1
g114037.t1
g72755.t1
g165975.t1
g47561.t1
g6132.t1
g154192.t1
g90475.t1
g62193.t1
g170389.t1
g145949.t1
g69675.t1
g152608.t1
g107955.t1
g179563.t1

**Supplementary Figure 3. Phylogenetic tree of duplicated genes.**

Phylogenetic tree of (a) Catalase, (b) SOD, (c) GST, and (d) HSP. These phylogenetic trees contain each gene of *R. varieornatus* and *C. elegans*. Multiple alignments were conducted using MAFFT and phylogenetic trees were constructed by FastTree. Bootstraps are showed under the branch. Red lines indicate highly similar orthologs (> 99% identity).

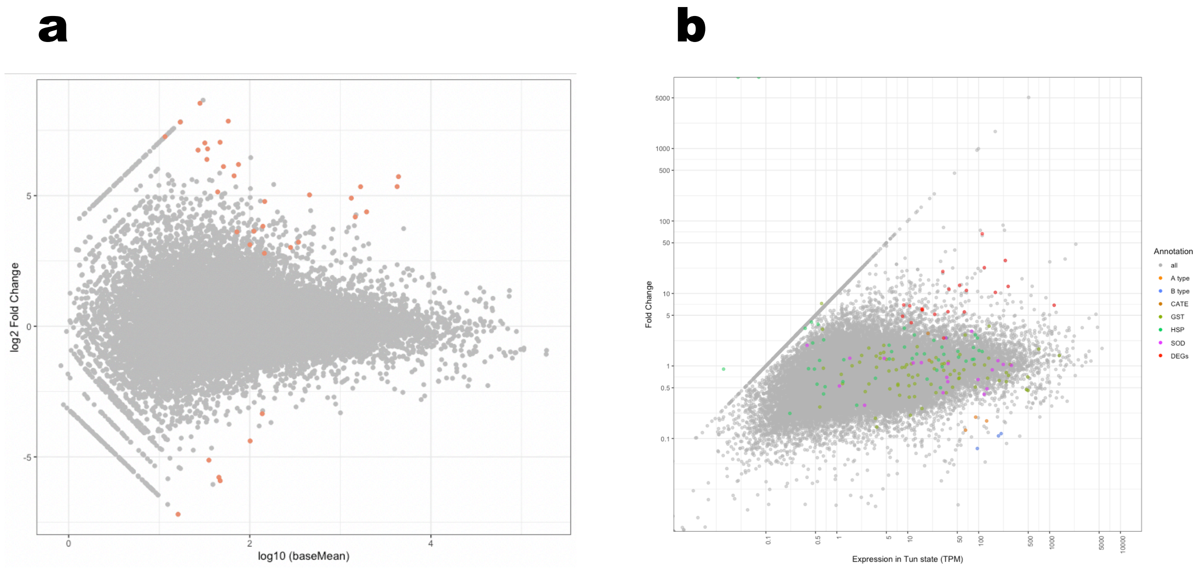


**Supplementary Figure 4. M-A plot of DEGs and plot of fold change-gene expression in anhydrobiosis state.**

(a) Data represent individual genes and plotted by gray. FRD < 0.05 were defined as DEGs and colored in the red plot. The vertical axis shows log2 fold-change and the horizontal axis shows log10 baseMean, with positive change indicating up-regulated genes and a negative change indicating ting the down-regulated genes. (b) All genes were plotted in gray and genes that likely contribute to anhydrobiosis were colored. The vertical axis shows expression level (Transcript per million; TPM) in anhydrobiosis state and the horizontal axis shows fold change between active and anhydrobiosis state.

### Supplementary Tables

| Genome type | | Original | Final (max TPM>1 cut off) |
|---|---|---|---|
| BUSCO (genome) (%) | Complete | 89.4 | 87.8 |
| | Single copy | 40.9 | 39.6 |
| | Duplicated | 48.5 | 48.2 |
| | Flagmented | 3.6 | 4.6 |
| | Missing | 7.0 | 7.6 |
| BUSCO (augustus.aa) (%) | Complete | 93.7 | 92.7 |
| | Single copy | 23.1 | 25.4 |
| | Duplicated | 70.6 | 67.3 |
| | Flagmented | 3.0 | 3.0 |
| | Missing | 3.3 | 4.3 |
| Scaffold number | | 107,804 | 30,095 |
| N50 (bp) | | 4891 (#22099) | 6674 (#7409) |
| Total length (bp) | | 388,783,196 | 153,718,311 |
| Gene number | | 186,205 | 42,608 |

**Supplementary Table 1. Screening of genome assembly.**

**a**

| All reads | Mapped | Unmapped | Mapping promotion |
|---|---|---|---|
| 49,410,721 | 41,640,594 | 7,770,127 | 84.27% |

**b**

| | Mapped | Unmapped | Mapping promotion |
|---|---|---|---|
| act1 | 18,375,873 | 6,443,782 | 74.04% |
| act2 | 26,119,645 | 10,371,933 | 71.58% |
| act3 | 24376861 | 8,760,852 | 73.56% |
| tun1 | 12,625,019 | 4,825,469 | 72.35% |
| tun2 | 13,636,144 | 4,737,143 | 74.22% |
| tun3 | 17,533,800 | 6,769,570 | 72.15% |

**Supplementary Table 2. Mapping ration of DNA-Seq and RNA-Seq.**

(a) Mapping proportions of DNA-Seq data (against BRAKER assembled genome). (b) Mapping proportions of RNA-Seq data (against BRAKER assembled genome using TopHat2)

### Supplementary Tables

| Transcriptome Statistics | Original | Final |
|---|---|---|
| Total transcripts | 147,063 | 20,704 |
| Components (seq) | 283,876 | 63,930 |
| GC contents (%) | 46.76 | 46.26 |
| Total length | 231,346,032 | 83,283,603 |
| N50 | 1,364 (#47,853) | 1,955 (#13,084) |

**Supplementary Table 3. Overview of transcriptome assembly.**

**Supplementary Table 4-7are supplied as Supplementary Data.**
